# Temporal dynamics of ectomycorrhizal fungi: Leaf habit and exploration strategy contribute to seasonal variation in community abundance and composition

**DOI:** 10.1101/2025.06.20.660784

**Authors:** Nicholas Medina, Kelsey Patrick, Teodora Nikitin, Claire Kaliski, Aubrie Bogle, Marvin Lo, Peter G. Kennedy, M. Luke McCormack

## Abstract

Ectomycorrhizal (EcM) fungi are well-recognized symbionts impacting tree health and ecosystem functioning globally, yet understanding of their timing of proliferation in soils across seasons and years remains limited. We analyzed monthly patterns of EcM fungal abundance and community structure over two years in five temperate monodominant forest plots via quantitative PCR and Illumina sequencing. We found that the phenological dynamics of EcM fungi differed significantly by host tree leaf habit, fungal exploration type, fungal genus, and soil moisture. Overall, total EcM fungal abundances based on qPCR consistently peaked in autumn, and were more dynamic in evergreen than deciduous plots, supporting ideas of surplus carbon and asymmetric above-belowground dynamics. Longer-distance exploration types peaked earlier and were more stable than shorter-distance types, suggesting an independent and supportive role in releasing spring nutrients. About half of 20 focal taxa consistently peaked in either autumn, summer, or spring, while others were either host- and/or year-dependent. Our findings highlight that phenology is a key EcM fungal trait best explained by both host and fungal contributions, and future studies across biomes should consider seasonal shifts and sampling to elucidate phenological traits.

**Summary:**

- The timing of belowground production and seasonal community dynamics remain poorly understood for ectomycorrhizal (EcM) fungi.
- We collected soils monthly for two years from five temperate monodominant forest plots.
- Fungal production peaked in autumn, shorter-distance and evergreen-associated spanned wider ranges, and half of focal fungal genera showed seasonal preference, emphasizing autumn surplus carbon and spring nutrients from long-distance types.
- Future studies should consider seasonal shifts when sampling EcM fungal communities, and forest carbon models should include asymmetric above-belowground phenology.

**Translated Summary (Spanish):**

- La fenología de la producción y composición de comunidades de hongos ectomicorrízicos (EcM) es poco estudiada.
- Recolectamos suelos mensualmente por dos años de cinco parcelas mono-dominantes templados.
- Producción máxima de hongos ocurrió en otoño, hongos asociados con árboles siempreverdes y de exploración de corta-distancia observaron rangos más amplios, y la mitad de géneros de hongos focales observaron preferencia estacional, enfatizando extra carbono en otoño y nutrientes en primavera de tipos larga-distancia.
- Estudios deben considerar cambios estacionales para el muestreo de hongos EcM, y modelos de carbono deben incluir fenología asimétrica entre hojas y hongos.

**Plain language summary:** Ectomycorrhizal fungi are critical for the global carbon cycle, but their seasonal and inter-annual growth patterns remain unclear. We sample soil DNA monthly over two years across five different monodominant temperate forest stands. We find an overall belowground peak in autumn, with significantly later growth under wetter conditions, more dynamism with evergreen trees, and distinct spring growth by longer-distance fungi.

## Introduction

The timing of annual biological growth and events in relation to abiotic and biotic conditions, or phenology, is a fundamental attribute of ecosystems, especially in seasonal biomes (Cleland *et al*., 2007). Long-term phenological changes reflect ongoing climate changes, with either earlier spring (Meng *et al*., 2020) or species-dependent responses (Kauserud *et al*., 2012; Tuno *et al*., 2020), with potentially important cascading effects on consumer-resource mismatches (Liu *et al*., 2022) and ecosystem resilience to stress factors such as to drought (Li *et al*., 2023). To date, however, most phenology studies focus on aboveground components of ecosystems (Flint, 1974), such as bud break (Ware *et al*., 2021), reproduction, or senescence (Dox *et al*., 2020), often in leaves but also including aboveground structures of forest floor activity such as mushrooms (Kauserud *et al*., 2008; Tuno *et al*., 2020; Michaud *et al*., 2024) and culturable decomposers (Katz & Lieth, 1974). The abundance of aboveground phenology data coupled with limited belowground data often leads Earth system model structures to assume above-belowground links (McCormack *et al*., 2012; Nakahata *et al*., 2021), including assumptions regarding symmetry and synchrony of growth patterns, although recent studies increasingly suggest asymmetric above-belowground phenology patterns (Steinaker *et al*., 2010; Abramoff & Finzi, 2015; Michaud *et al*., 2024). Indeed, models that explicitly represent belowground microbial processes yield more realistic predictions, such as to responses to warming (Wieder *et al*., 2013), highlighting the importance of adding belowground phenology into models based on empirical data, particularly of distinct fine-root pools and microbial symbionts such as mycorrhizal fungi (Wang *et al*., 2023). To do this, we need to better understand the fundamental empirical phenology traits of belowground components (Radville *et al*., 2016; Chaudhary *et al*., 2020; Schaffer-Morrison & Zak, 2023).

The phenology of fine roots is a key mediator of energy inputs and microbial activity across ecosystems. Several relevant studies point to peak root production in summer and autumn, but with notable exceptions (King *et al*., 2002; McCormack *et al*., 2014a, 2015) and inter-annual variation (Mariën *et al*., 2021). Fundamentally, root growth is thought to be driven by soil temperature and moisture, with seasonal (Nakahata, 2022) and inter-annual variation in influence (Withington *et al*., 2021). Root growth can also be driven by leaf habit, where the potential for year-round photosynthesis allows evergreen trees to rely less on belowground storage tissues, and potentially yield more available and excess photosynthates (Prescott *et al*., 2020), compared to deciduous trees (Bourdeau, 1959; Warren & Adams, 2004). Indeed, available data on the phenology of non-structural carbohydrates show a relatively more uniform amount of non-structural carbohydrates throughout the year in an evergreen species compared with a strong peak in autumn for deciduous species (Furze *et al*., 2019). Drivers of root phenology may then shape the phenology of rhizosphere symbionts (McCormack *et al*., 2015), including mycorrhizal fungi, which are under-recognized drivers of global carbon storage and ecosystem functions (Peay *et al*., 2016; Peay, 2016; Hawkins *et al*., 2023).

The phenology of ectomycorrhizal (EcM) fungal production is likely controlled by both host and fungal contributions, including their individual responses to soil microclimate, fine-root inputs and host tree preferences (Dickie *et al*., 2010; Tedersoo *et al*., 2024), above-belowground energy tradeoffs by host trees (Michaud *et al*., 2024), and temporal niche partitioning by fungal species (Koide *et al*., 2007; Courty *et al*., 2008; Pickles *et al*., 2010). Seasonal patterns in fine root production and activity patterns point generally to late summer and autumn peaks in EcM fungal phenology (McCormack *et al*., 2014b; Withington *et al*., 2021). Autumn peaks also align with when surplus C from photosynthesis is produced, which supports emerging ideas that surplus C drives rhizosphere microbiome dynamics (Prescott *et al*., 2020; Bunn *et al*., 2024; McCulloch *et al*., 2026). Additional phenological variation may emerge from host differences in physiology, where longer photosynthesis windows may allow more dynamic growth by helping alleviate soil microbial energy limitations (Cotrufo et al., 2013; Hoehler & Jørgensen, 2013; Gunina & Kuzyakov, 2022). EcM fungal phenology may also vary by fungal traits, particularly different hyphal exploration types, which range from contact and short-distance extramatrical hyphae emanating from the host root, to medium- and long-distance types exploring up to meters (Agerer, 2001). These traits likely contain additional under-studied variation (Jörgensen *et al*., 2023) yet nonetheless still have important implications for enzyme activity (Courty *et al*., 2007, 2010). Exploration types may differ in their phenology, as longer-distance types such as *Suillus* (Lofgren *et al*., 2024) may need more resources (carbon and/or nutrients) to construct their tissues compared to contact types such as *Russula* (Ekblad *et al*., 2013). Such different resource requirements, along with potentially more soil- and detritus-directed saprotrophic enzyme abilities (Koide *et al*., 2008; Smith *et al*., 2017), may lead longer-distance types to show less variable phenology, and even earlier growth during spring host tree activity, compared to contact types. In contrast, contact types may rely more on root inputs, leading them to more closely match root growth in autumn, when excess photosynthates may be allocated belowground (Prescott *et al*., 2020; Bunn *et al*., 2024). As a result, surplus C in autumn should predict stronger autumn peaks in contact types compared to longer-distance types with some independent saprotrophic capacity (Koide *et al*., 2008). Alternatively, higher growth earlier in spring may reflect microbial immobilization of otherwise leachable nutrients, supporting the vernal dam hypothesis (Bormann *et al*., 1977; Rothstein, 2000). Overall, fine-resolution patterns in EcM phenology remain under-studied.

To better document patterns of belowground EcM fungal phenology, this study analyzes the belowground seasonality of EcM fungi across a diverse set of temperate host tree species that differ in functional traits. We addressed the following questions: 1) Is total EcM fungal belowground production seasonality driven by soil moisture and temperature; 2) how do these patterns vary across host tree species, specifically evergreen versus deciduous trees; and 3) how does seasonality vary among EcM fungal species and functional groups? We hypothesized that: 1) EcM fungal production would be significantly driven by soil microclimate (i.e., moisture and temperature), similar to sporocarp responses (Kauserud *et al*., 2012); 2) EcM fungal production would be more dynamic under deciduous host trees compared to evergreen trees, due to more variation in nutrients from more concentrated periods of leaf senescence; and 3) EcM fungal phenology would vary by hyphal exploration types, due to differences in tissue construction, as well as by fungal genera given host tree species preferences (Tedersoo *et al*., 2024). Finally, we synthesize our results and consider their alignment with two different perspectives for growth and allocation in forest ecosystems, namely the C surplus C hypothesis (Prescott *et al*., 2020) which would be best supported by greater EcM fungal abundance later in the growing season, compared with the vernal dam hypothesis (Bormann *et al*., 1977; Rothstein, 2000) which would be more consistent with higher EcM fungal growth earlier in the growing season.

## Methods

### Study site

This study was conducted using five monodominant forestry plots at the Morton Arboretum in Lisle, Illinois, USA (41.81 N, 88.05 W), further described by Midgley and Sims (2020). Briefly, the forestry plots resemble a common garden experiment, with similar soil chemistry across primarily Alfisol and some Mollisol soil orders (Midgley & Sims, 2020). Annual air temperature ranges 7-29 °C and mean annual precipitation is near 1000 mm since the year 2008. Soil moisture and temperature data were collected at the site level using HOBO data loggers (Onset Computer Corporation, Bourne, MA, USA) installed at 10 cm depth in the topsoil, and these data were very similar to plot-level measurements available in a subset of plots. Plot host tree species used were: *Carya ovata* (shagbark hickory), *Quercus alba* (white oak), *Quercus bicolor* (swamp white oak), *Picea abies* (Norway spruce), and *Pinus strobus* (Eastern white pine). All five plots were of similar stand ages and management, and were on Alfisols soils, except *Q. bicolor*, which was on a Mollisol. Each plot had a distinct leaf habit (i.e., evergreen or deciduous), which also aligned with plant group (i.e., gymnosperm or angiosperm, respectively).

### Soil sampling

In each plot, soils were sampled monthly from six fixed 30 × 160 cm subplots that were 10 m apart on average and cored down to 10 cm depth using an autoclaved round 1 cm-wide metal tube. In each subplot, monthly samples were spaced 20 cm apart along a transect spanning the subplot’s long side, and annual transects were adjacent. Soil cores were stored in a cooler until the soil was taken out, homogenized, and subsampled later the same day. Soil subsamples were stored in 2 mL screw-cap vials at -80 °C for up to six months until lyophilization, which was done at once for all samples within a year and stored in a dark cabinet at room temperature within two weeks of DNA extraction, which then occurred over two months.

### DNA extraction, quantitative PCR, and sequencing

Approximately 0.25 g of soil was subsampled and used for DNA extraction with Qiagen DNeasy PowerSoil Pro kits following the manufacturer’s protocol, but homogenized for 30 sec at 4000 rpm using a MP FastPrep tissue homogenizer, both before weighing and after the bead beating step. Extracted DNA was stored at -80 °C for approximately two months until quantitative PCR (qPCR) was done to approximate fungal biomass, and library preparation according to Gohl et al. (2016), followed by sequencing of the ITS2 region using 5.8SR (5’ TCGATGAAGAACGCAGCG 3’) and ITS4 (5’ TCCTCCGCTTATTGATATGC 3’) primers on the Illumina NextSeq platform (V3 chemistry) at the University of Minnesota Genomics Center.

### Bioinformatics

Raw sequence files were initially processed with cutadapt (Martin, 2011) to identify all sequence reads with primers and sequencing adapters and remove both. The remaining sequences were filtered, denoised, and merged using the DADA2 algorithm (Callahan *et al*., 2016) using recommended parameters for fungal ITS (*maxN* = 0, *maxEE* = c(2, 2), *truncQ* = 2, *minLen* = 50). Amplicon sequence variants (ASVs) were assigned taxonomy using the UNITE (v9, Abarenkov et al., 2024) database. All forward and reverse .fastq files were deposited in the NCBI Short Read Archive under BioProject ID PRJNA1293086. ASVs allow greater taxonomic resolution but with similar ecological results compared to traditional operational taxonomic units (OTUs) based on 97% sequence similarity (Glassman & Martiny, 2018). Following general recommendations for high-throughput sequencing (Lindahl *et al*., 2013) and account for possible index bleed (Carlsen *et al*., 2012), all ASVs with fewer than ten reads were eliminated. All remaining ASVs were assigned exploration types using the FungalTraits database (Põlme *et al*., 2020), acknowledging some categorical flexibility (Jörgensen *et al*., 2023; Johnson & Marín, 2025).

### Statistics

All analyses were performed using R version 4.4 (R Core Team 2025). Site-level soil temperature and moisture data were smoothed using rolling averages from the previous week and filtered for those values on sampling dates. To obtain total EcM fungal abundance, total fungal abundance values from qPCR were multiplied by a factor of 0.17, which was equal to the proportion of sequenced reads matching to EcM lifestyles in the ITS2 community data and stayed consistent over time. This approach has been used in multiple recent studies to provide guild-specific abundance estimates (Mielke *et al*., 2025; Boeraeve *et al*., 2025). To compare fungal communities across samples, we rarefied to a common lowest number of observed sequence reads in each year (approx. 50,000 and 30,000 reads), which excluded five and three (<2%) samples, as rarefaction remains appropriate for microbiome data (Schloss, 2024). Genus-level and exploration type level abundances were summed across ASVs. Genus- and group-level data were centered across subplots within a forest plot using median values. Visualizations of the results show zeros adjusted by adding one if log-transformed.

Hierarchical generalized additive mixed models (GAMMs) were used to analyze time series data (Pedersen *et al*., 2019). To prepare response metrics, EcM fungal abundances based on qPCR were transformed to proportions of subplot maximums within each year to account for batch effects from yearly sequencing. Rarefied trait- and genus-level abundances were summed across ASVs within a sample, and maximum annual values calculated for exploration types to best match raw data. Twenty focal genera were chosen for analysis based on high abundance in our data across plots and frequent discussion in the literature. Values that were NA were omitted during model runs and ordinations as necessary as well as zero values adjusted by one to improve model convergence. Distribution families were tweedie *tw()* or negative binomial *nb()*, allowing automatic parameter fitting available with *mgcv* but not *gamm4*, chosen based on best fit to model assumptions shown by the *appraise()* function in the *gratia* package (Simpson, 2024). Model structure included a global smoother for unique week number between years to distinguish samples within a calendar month, with maximum curvature knots set to *k*=17 given 19 total sampling times, plus a similar smoother separated by trait or genus using the *by* argument, and *m*=1 in the global week smoother to reduce expected concurvity. Soil temperature and moisture were included as separate *ti()* terms with a *ti()* interaction. Fixed effects included leaf habit and random effect *bs=‘re’* was subplot number up to six, with plot-level variation considered qualitatively to avoid model overfitting. The model *method = ‘fREML’* and *mgcv::bam()* was needed to reduce runtime. After modeling, we assigned each focal genus a seasonal preference if, after filtering for presence in at least three samples per year, it peaked in the same season each year in over half of the plots where it was detected (e.g. at least three, or one if specialist), and instead assigned host- or year-dependent if peak season varied either between years or across plots.

For the community analyses, both years of sequence run data were combined into a genus-level community matrix with summed and square-root (Hellinger) transformed abundances (Legendre & Gallagher, 2001) with *vegan::decostand()* to maintain influences of dominant taxa compared to center-log-ratio (Aitchison) transformation (Gloor *et al*., 2017; Fuschi *et al*., 2025). Genus-level community analysis facilitated generalizable discussion and ordination, resolved separate ASV identifiers per year, and showed similar statistical results to using the species-level abundance matrix, with some variation in vector direction for abundant species (Supplementary Fig. S3, Table S7). Differences in centroids by main study factors were tested with a PERMANOVA based on Bray-Curtis distances and 999 default permutations using the *adonis2()* function from the *vegan* package (Oksanen *et al*., 2001; Dixon, 2003). To parse between centrality differences and variance in PERMANOVA (Anderson, 2001), we also explored community dispersion as a measure of beta diversity (Anderson, 2006; Anderson *et al*., 2006) with a PERMDISP using the *vegan*::*betadisper()* and *vegan::permutest()* functions, with a separate test for week and leaf habit due to *group* argument limitations. For community ordination linking linear environmental and species vectors, a redundancy analysis was done using the *vegan*::*rda()* function on the full community table including covariates followed by an *anova()* summary (Ramette, 2007).

## Results

### Sequencing

The quality-filtered final dataset included approx. 11 million sequence reads and 10 thousand unique ASVs. A total of 1,048 ASVs were assigned to the ectomycorrhizal lifestyle across years, representing approximately 17% of total reads. Among the other lifestyles assigned, the remaining majority were roughly equal proportions of saprotrophs (25%) as well as unknown lifestyles (25%). The majority (91%) of ASVs were assigned taxonomy at the genus level, with all those assigned ectomycorrhizal being mapped to a genus. Approximately 55% of the EcM fungal ASVs were at the species level, and a similar proportion of species were assigned across all other fungi detected (50%).

### Total EcM fungal abundance based on qPCR

EcM fungal abundance changed significantly and consistently over time by host tree leaf habit (*dev. expl*. = 20.4%) (Supplementary Table S1), showing annual unimodality (Fig. 1a), and abundance showed wider ranges in evergreen (*EDF* = 3.3, *F* = 13.7, *P* < 0.0001) compared to deciduous plots (*EDF* = 1.5, *F* = 18.9, *P* < 0.0001). EcM fungal abundance was consistently lowest by the spring equinox at week 12 across plots and years. Peaks in abundance were consistent across plots and their leaf habit, but production varied by year (*est*. = -1.2 ± 0.2, *t* = - 7.3, *P* < 0.0001), showing peaks by the autumn equinox around week 38 in the first year, and peaks by the winter solstice around week 50 in the second year. The range in EcM fungal abundance was smallest in abundance in the *C. ovata* plot, although changes in EcM fungal abundance were not explained by observed temperature or moisture patterns, as their interaction remained non-significant (*EDF* = 1, *F* = 2.7, *P* = 0.10) (Fig. 1b).

**Fig. 1.**
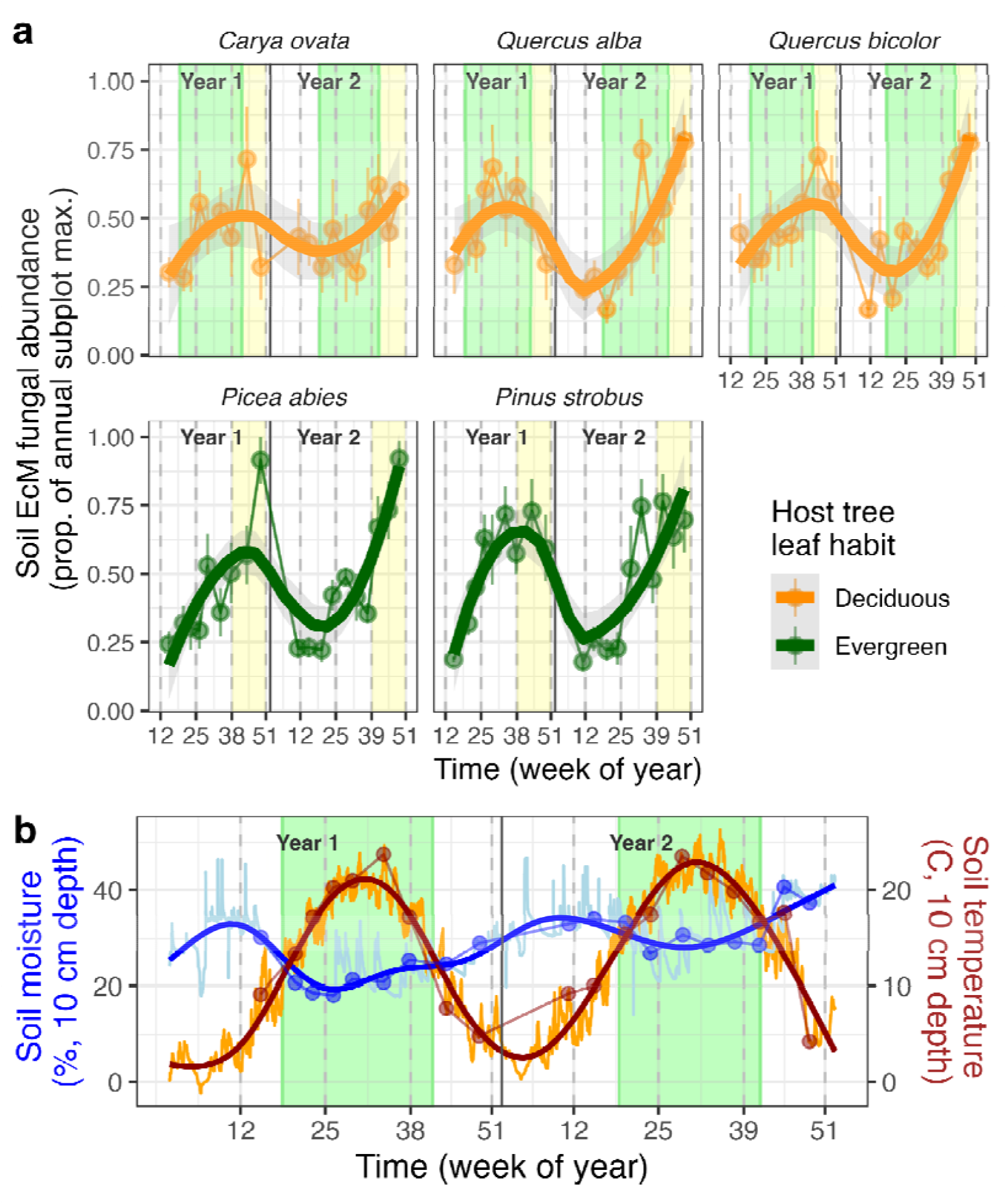
Total abundance of soil EcM fungi peaks in autumn. (**a**) EcM fungal abundances in soils over time across five temperate monodominant forest plots, with deciduous (top row) and evergreen (bottom row) leaf habits. Points show mean ± 1 standard error (SE) of subplot-level proportion of annual subplot maximum values (n = 6), across approximately monthly sampling times for 9 total in the year 2023 and 10 in 2024, and lines connect medians and smooth ‘loess’ curves (n = 14, k = 17). Week of year number labels on x-axis indicate approximate equinox and solstice weeks, with the growing season window highlighted in green, and the autumn season is highlighted in yellow to highlight typical peak production. (**b**) Site-level soil moisture (left, blue) and temperature (right, orange) at 0-10 cm depth, with sampling dates as larger dots and solstices and equinoxes highlighted (grey dashed lines). Data shown include raw moisture (light blue line) with connected medians (thin blue line) and ‘loess’ smooth (thick blue line), as well as raw temperature (orange) with connected medians (thin red line) and ‘loess’ smooth (thick red line).

### EcM fungal exploration type

EcM fungal abundance changed significantly across seasons by exploration type (*Dev. expl*. = 13.8%), and also by year (*est*. = -1.9 ± 0.3, *t* = -5.9, *P* < 0.0001) (Fig. 2, Supplementary Table S2). Specifically, the abundance of long-distance types changed noticeably over time, albeit relatively stably (*EDF* = 7.1, *F* = 11.3, *P* < 0.0001), whereas in contrast, contact type abundance was more dynamic (*EDF* = 4.2, *F* = 5.5, *P* < 0.0001), and also differed by host tree leaf habit. Short-coarse (*EDF* = 5.3, *F* = 7.9, *P* < 0.0001) and medium-smooth types (*EDF* = 1, *F* = 14.7, *P* = 0.0001) also varied widely and significantly across the years, but unlike contact types, these abundances instead stayed consistent between host tree leaf habits. All peaks and minima for dynamic exploration types matched overall EcM fungal production patterns, with similar peaks near autumn/winter weeks 38/50 and minima near spring equinox week 12, and with no separate or interactive influence of temperature. However, exploration type phenology was significantly affected by soil moisture variation (*EDF* = 7.1, *F* = 2.5, *P* = 0.008). Notably, medium-fringe (*EDF* = 6.3, *F* = 5.3, *P* < 0.0001) and short-delicate types (*EDF* = 1, *F* = 9.6, *P* = 0.0019) showed different patterns by host tree leaf habit, with short-delicates only variable over time in deciduous plots, and medium-fringe only variable over time in evergreen plots.

**Fig. 2.**
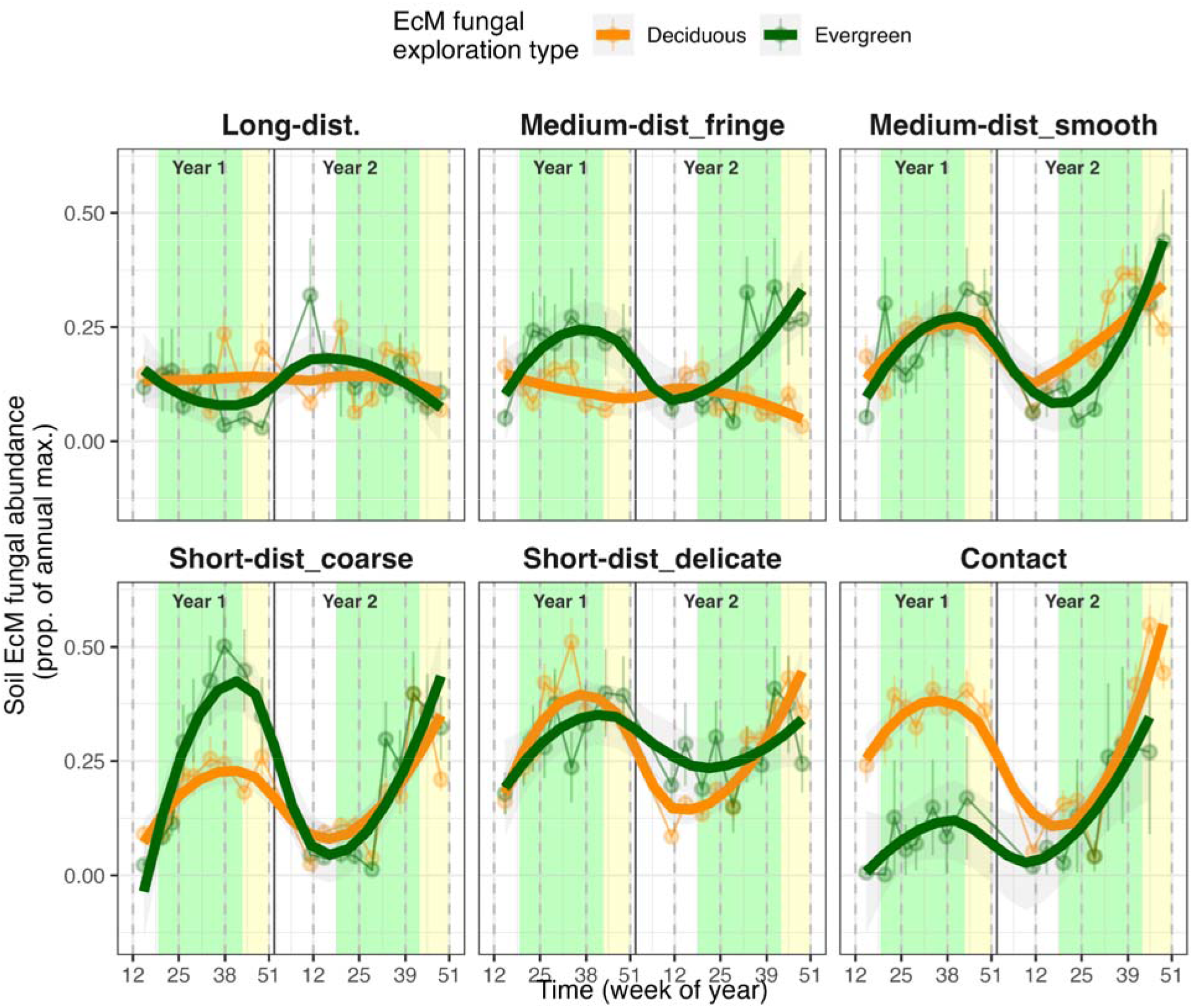
Shorter-distance EcM fungal exploration types are more seasonally dynamic than longer-distance types. Seasonality of ectomycorrhizal exploration types among five temperate monodominant forest plots, in either evergreen (dark green) or deciduous (orange) plots. Lines show smooth loess curves (n = 6) across mean values among proportions of annual maxima across subplots ± 1 SE for each of the 19 monthly sampling dates across the years 2023-2024 (k = 17). Week of year number labels on x-axis indicate approximate equinox and solstice weeks, with aboveground growing season windows highlighted in green, and the autumn season highlighted in yellow to highlight typical peak production.

### EcM fungal genera

The abundances of all 20 focal genera changed significantly across seasons (*Dev. expl*. 25.4%), with additional effects of host tree leaf habit (*est*. = 0.51 ± 0.06, *t* = 9.4, *P* < 0.0001), and year (*est*. = -2.8 ± 0.3, *t* = -10.6, *P* < 0.0001), in part driven by significant soil moisture changes (*EDF* = 2.2, *F* = 4.4, *P* = 0.045) and subplot variation (*EDF* = 4.8, *F* = 18.4, *P* < 0.0001) (Fig. 3, Supplementary Table S3). All genera showed unimodal peaks, but with different timing based on genus, host tree, and/or year (Fig 3b). Genera with spring (*ca*. week 18) preferences across both years included *Suillus, Rhizopogon*, and *Scleroderma* (*P* < 0.0001), while *Helvella* and *Thelephora* preferred summer (peaking *ca*. week 31) (*P* < 0.0001). Genera with consistent autumn (*ca*. week 40) peaks included *Tuber, Tomentella, Russula, Inocybe, Humaria*, and *Elaphomyces* (*P* = 0.0005). Several other genera varied in peak times by either host tree and/or year, including: *Wilcoxina, Sebacina, Lactarius, Hymenogaster, Hygrophorus, Cortinarius, Cenococcum*, and *Amanita* (*P* < 0.03), while *Piloderma* which did not show significant changes over time.

**Fig. 3.**
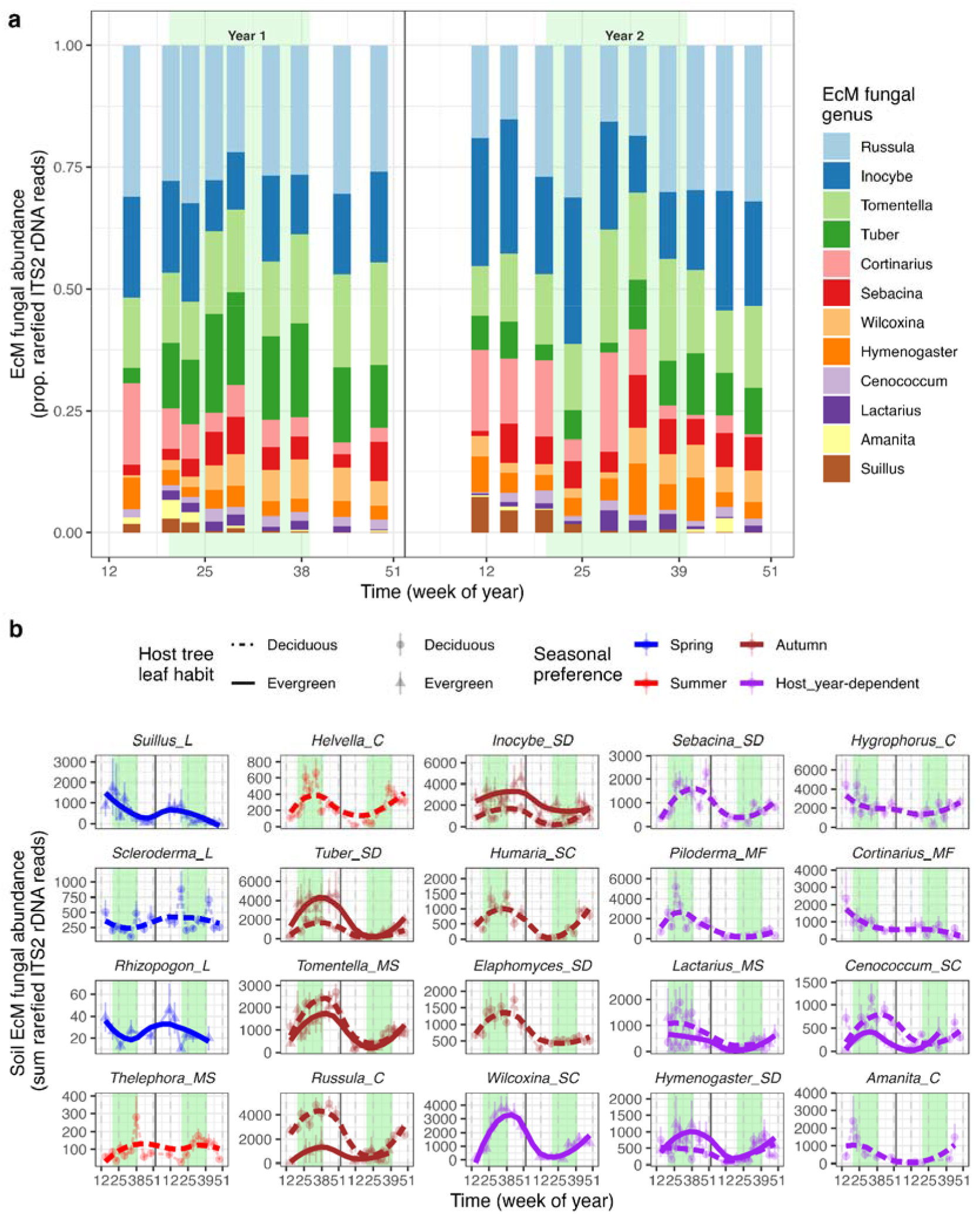
Abundant EcM fungi showed seasonal growth preferences. (**a**) Proportional compositional changes of top 10 soil ectomycorrhizal fungal genera across all six subplots and five plots over the two-year course of monthly sampling in this study. (**b**) Lines show seasonal abundance of each ectomycorrhizal fungal genus among five temperate monodominant forest plots, in either evergreen (triangle, solid line) or deciduous (circle, dashed line) host tree species plots, and colored by peak season spring (blue), summer (dark red), autumn (orange), or host or year-dependent (dark green). One line is shown in cases where the genus was clearly primarily found in that stand type. Lines show smooth loess curves (n = 6) across mean values among subplots ± 1 SE for the 19 total monthly sampling dates in the years 2023-2024. Week of year number labels on x-axis indicate approximate equinox and solstice weeks, the growing season window is highlighted in green.

### EcM fungal community composition

The composition of EcM fungal communities changed significantly over time (Fig. 4, Supplementary Table S4). Significant factors shaping community centroids included mainly host tree leaf habit (*DF* = 1, *F* = 176.8, *P* = 0.001, *R*^*2*^ = 0.238) (Fig. 4a,c), followed by week of year (*DF* = 1, *F* = 3.6, *P* = 0.002) and both soil moisture (*DF* = 1, *F* = 3.0, *P* = 0.01, *R*^*2*^ = 0.004) and temperature (*DF* = 1, *F* = 2.8, *P* = 0.007, *R*^*2*^ = 0.004) (Fig. 4c). Community clusters showed equal variance around centroids when separated by week and host tree leaf habit (Fig. 4a,d, Supplementary Table S5), implying similar overall shifts in relative abundances across shared genera (Fig. 4a,b,d). Significant environmental vectors mapping onto community ordination included all main ones tested, namely week (*DF* = 1, *F* = 1.8, *P* = 0.045), host tree leaf habit (*DF* = 1, *F* = 119.7, *P* = 0.001), and temperature (*DF* = 1, *F* = 2.4, *P* = 0.015), excluding soil moisture and any interactions (Fig. 4c, Table S6). While communities showed similar amounts of change, the directions of change differed over time between years and leaf habits (Fig. 4b). For instance, EcM fungal communities in deciduous plots moved counter-clockwise in ordination space in the first year, shifting from *Tuber* to *Inocybe* directions (Fig. 4d), and then instead in the opposite clockwise direction in the second year (see Fig. 3a). Conversely, EcM fungal communities in evergreen plots showed opposite patterns in both years compared to deciduous communities.

**Fig. 4.**
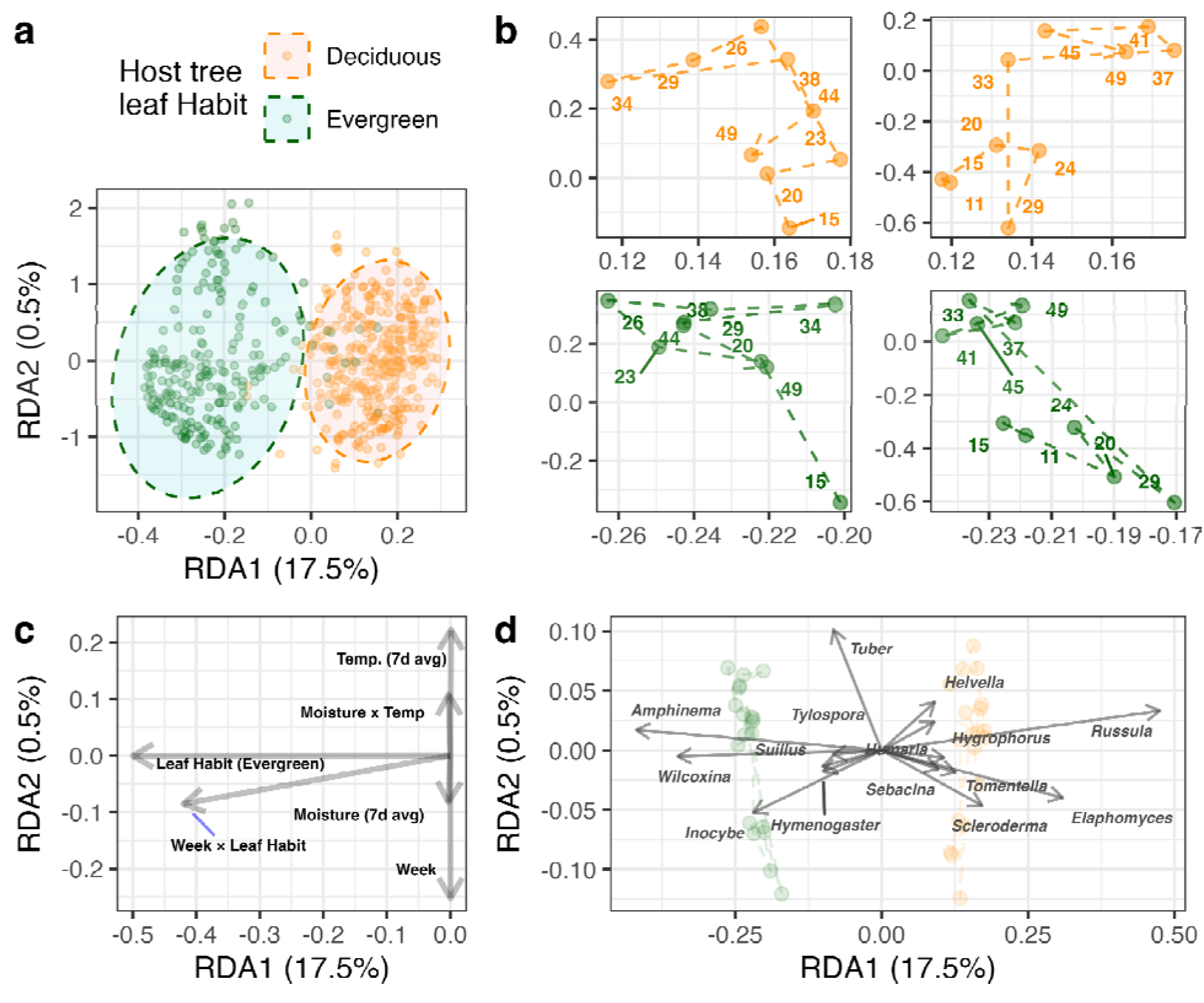
Changes in soil EcM fungal community composition were explained by week of year, leaf habit, soil temperature, and soil moisture. Ordination RDA on EcM fungal communities over two years across five monodominant temperate host tree stands. (**a**) Centroids and 95% confidence ellipses grouped by evergreen (green) and deciduous (orange) host tree leaf habits. (**b**) Phase-portrait trajectories of week-level centroids and week number labels for deciduous (orange, top row) and evergreen (green, bottom row) plots in years one (left column) and two (right column). (**c**) Scaled environmental and (**d**) vector arrows in the same RDA ordination space, with a transparent version of panel **b** placed under panel **d**.

## Discussion

Our results support the hypotheses that EcM fungal seasonality is strongly driven by both host and fungal contributions, including fungal functional group, fungal taxonomy, host tree species and leaf habit, and to a lesser extent, climatic factors such as soil moisture, similar to sporocarp phenology and in support of our first hypothesis. In contrast to our second hypothesis, our findings show more dynamic seasonal patterns in evergreen versus deciduous host tree species plots, and in line with our third hypothesis, we also observed more temporal variation among shorter-distance exploration types compared to longer-distance types. The greater frequency of autumn over spring peaks for EcM genera also helps distinguish among plant-versus soil-based drivers of EcM fungal growth. Specifically, the more common autumn peaks in belowground growth of most EcM fungi are consistent with responding to surplus host C (Prescott *et al*., 2020), with some exceptions for spring growth of long-distance exploration type taxa that may align with nutrient resorption predictions of the vernal dam hypothesis instead (Bormann *et al*., 1977; Rothstein, 2000).

The difference in EcM fungal seasonality by host tree leaf habit aligns with differences in carbon budget dynamics between deciduous and evergreen trees. Although deciduous trees tend to show more distinct seasonal patterns of carbohydrate allocation pulses to leaves in spring (Krishna & Chandra Garkoti, 2022) and more variability in root exudate and non-structural carbohydrate production than evergreen trees (Furze, 2018; Furze *et al*., 2019), this was not mirrored in belowground EcM fungal growth. Instead, we observed more dynamic EcM fungal growth under evergreen plots, suggesting that EcM fungal growth overall, including of several genera and for several key short-distance exploration types, may be promoted by extra C availability from continuous photosynthesis, in part supporting the surplus C hypothesis (Prescott *et al*., 2020; Bunn *et al*., 2024). More dynamic EcM fungal growth under continuous photosynthesis also suggests above-belowground asymmetry in forest phenology and resource dynamics.

The more dynamic growth patterns of shorter-distance compared to longer-distance exploration types may reflect more responsiveness to host physiology and soil nutrient dynamics, in part due to lower tissue construction costs. Exploration types have been recently suggested to represent turnover (Jörgensen *et al*., 2023) instead of morphology (Agerer, 2001), where short-distance types have faster mycelial turnover and responses to nutrients, potentially favoring inorganic nutrients (Hobbie & Agerer, 2010) akin to broader mycorrhizal nutrient economies in forests (Phillips *et al*., 2013). The seasonality of short-distance exploration types (e.g., *Russula, Tuber, Cenococcum*) may be more closely associated with root physiology and seasonality, as non-structural carbohydrate dynamics are likely important for EcM symbioses (Sapes *et al*., 2021). This could also reflect autumn season preference of EcM fungi that was triggered by physiological cues of changes from dry and warm to wet and cool summer (Krah *et al*., 2023). In contrast, more stable production of longer-distance exploration types (e.g., *Suillus, Rhizopogon*) support their overall more likely responsiveness and/or tolerance of soil (e.g., drought or low-nutrient) conditions, as they tend to have more extramatrical hyphae and hydrophobic rhizomorphs (Parrent & Vilgalys, 2007; Lilleskov *et al*., 2011) with associated enzyme activity, which may still offer key functions even at relatively lower abundances (Hupperts *et al*., 2017). Within short-distance exploration taxa, short-coarse types showed greater range between mean minimum and maximum abundance under evergreens, and short-delicates varied more in deciduous host tree species plots. These then notably differ from long-distance and medium-distance fringe exploration types which displayed much less variation which overall emphasizes seasonality, and phenology more broadly, as a key distinguishing trait axis for mycorrhizas, with implications for soil functions (Argiroff *et al*., 2022; Schaffer-Morrison & Zak, 2023). Future work should continue characterizing seasonal patterns in finer-resolution EcM fungal-host tree combinations, and possible intransitive ecological connections (Soliveres & Allan, 2018; Soliveres *et al*., 2018) to help explain changing relative influences of seasonal drivers (Gallien *et al*., 2017).

Alongside exploration type, EcM fungi showed clear temporal separation by genus, in both year and host-dependent and independent ways. Importantly, multiple mechanisms may be at play throughout the year. Specifically, intrinsic growth strategies of different taxa and drought tolerance are likely important in spring and summer, while responsiveness to available host plant photosynthates are likely important in late summer and autumn. For example, the rhizomorphic growth habits of long-distance taxa such as *Suillus* and *Rhizopogon* may enable them to grow more readily in spring owing to mycelia or tissues that may already be established through longer-lived rhizomorphs (Pritchard *et al*., 2008; McCormack *et al*., 2010) persisting through winter, resembling advantages of successional priority effects (Kennedy *et al*., 2009, 2020) including on host establishment patterns (Policelli *et al*., 2019; Chase *et al*., 2020). Additionally, the potential use of organic matter degrading enzymes could enable them to grow early in spring even if active allocation of photosynthates belowground is limited due slowed canopy photosynthesis or competition with other plant carbon sinks in spring and early summer (e.g. leaves and wood growth). As soils warm and moisture declines over summer, the hydrophobic tendency of long-distance taxa may also provide an advantage if they are able to persist during drought conditions, and heat-responsive taxa may also be favored. However, as conditions change and if host plant carbon allocation belowground increases in late summer and autumn, other taxa may be favored instead. Specifically, shorter-distance genera may peak in autumn due to faster responses to timely root and carbohydrate pulses in cooler and wetter conditions, nutrient availability and decomposing enzyme production, and summer drought persistence, such as in speciose genera including *Russula* (Looney *et al*., 2018; Argiroff *et al*., 2022), *Tuber* (Ceccaroli et al., 2011), and *Tomentella* (Voříšková *et al*., 2014; Lu *et al*., 2022; Cheng *et al*., 2023). Overall, genus-level phenology patterns likely reflect understudied response traits, and possibly temporal niche partitioning (Koide *et al*., 2007), of key genera with various energetic and resource needs.

Several EcM fungal genera showed inter-annual and host-dependent variation, which revealed flexibility in some groups of EcM fungi. Mainly, the second wetter year likely pushed back summer peaks into autumn for nearly all focal general in this category, highlighting soil moisture as a significant and shared environmental cue broadly shaping EcM fungal phenology. For example, among those showing this inter-annual delay was *Sebacina*, which has shown higher abundances in both warm and wet conditions (Beidler *et al*., 2023). In addition to year-dependent peaks, some genera also showed host-dependent peaks, which was most clearly in short-coarse *Cenococcum*. Generally, *Cenococcum*’s phenology may be explained by integrated responses throughout a relatively long mycelial lifespan (Fernandez *et al*., 2013; McCormack *et al*., 2017), which may also lead to priority effects allowing successful primary or secondary establishment on new fine-roots. Finally, some genera may simply be very seasonally flexible and driven more strictly by cool and wet soil conditions, showing peaks in either spring and/or autumn, as seen in *Cortinarius, Hygrophorus*, and *Amanita* (Fig. 3b).

Our findings offer clear novel fine-scale resolution of EcM fungal seasonality, yet we also acknowledge some common data and inference limitations. For instance, using qPCR to approximate fungal biomass has precedent but is not unbiased (Mielke *et al*., 2025; Boeraeve *et al*., 2025), although alternatives such as hyphal ingrowth bags or ergosterol measurements also have their own unique tradeoffs (Wallander *et al*., 2013; Baldrian *et al*., 2013; Zuev *et al*., 2019). Recent advances in long-read sequencing may also offer finer taxonomic resolution to better parse species-specific seasonal patterns in species-rich EcM fungal genera (Tedersoo *et al*., 2021). Applicable to all soil DNA studies, it is also acknowledged that soil DNA can be persistent or relic (Carini *et al*., 2016), i.e., integrating past activity, so we recommend future work analyzing shorter-lived RNA to clarify transcriptionally active fungi, including potentially more rapid pulse changes (e.g., high rain after drought). Finally, longer-term similar datasets would clarify inter-annual response traits, as would additional studies across sites and forest types globally.

Overall, our results show significant seasonal variation in EcM fungal communities and relative abundances of dominant taxa, with important implications for tree and forest production dynamics. This was particularly true in evergreen host plots, which are typically thought to provide a more consistent and even C resource base for EcM fungi compared to deciduous hosts. Among exploration types, shorter-distance EcM fungi like *Russula* and *Inocybe* play a more important role in mediating belowground production dynamics in evergreen host plots, whereas longer-distance EcM fungi like *Suillus* and *Rhizopogon* remain more stable while associating with nutrient release during spring. Further, our results challenge current assumptions that EcM fungal production mirrors soil abiotic conditions or host tree production, given the notable additional variation among phenological groups of EcM fungi reported here (Wang *et al*., 2023). Given that many of our EcM fungal genera are globally common (Tedersoo *et al*., 2012), these patterns may also be generalized for similar ecosystems. Ultimately, our findings support recent calls asserting that mycorrhizal fungi warrant explicit and unique parameterization of their seasonality in Earth system models to improve predictions in response to global changes.

## Acknowledgements

We thank lab volunteer Bill Prescott for helping process soil sampling and equipment, as well as Ryo Nakahata and Kevin Li for statistical consultation. This project was funded by a US Department of Energy award #DE-SC0023480 to M.L. McCormack and a US National Science Foundation award #2129312 to P.G. Kennedy.

