## Supplementary material for "Temporal dynamics of ectomycorrhizal fungi: Leaf habit and exploration strategy contribute to seasonal variation in community abundance and composition": SI

### **Supporting Information / Online Appendix**

Structure below mirrors that of the main text (Greenbaum *et al.*, 2017). Statistical GAM output tables are organized and presented using the *‘flextable’* R package.

#### **Results**

##### Total EcM abundance based on qPCR

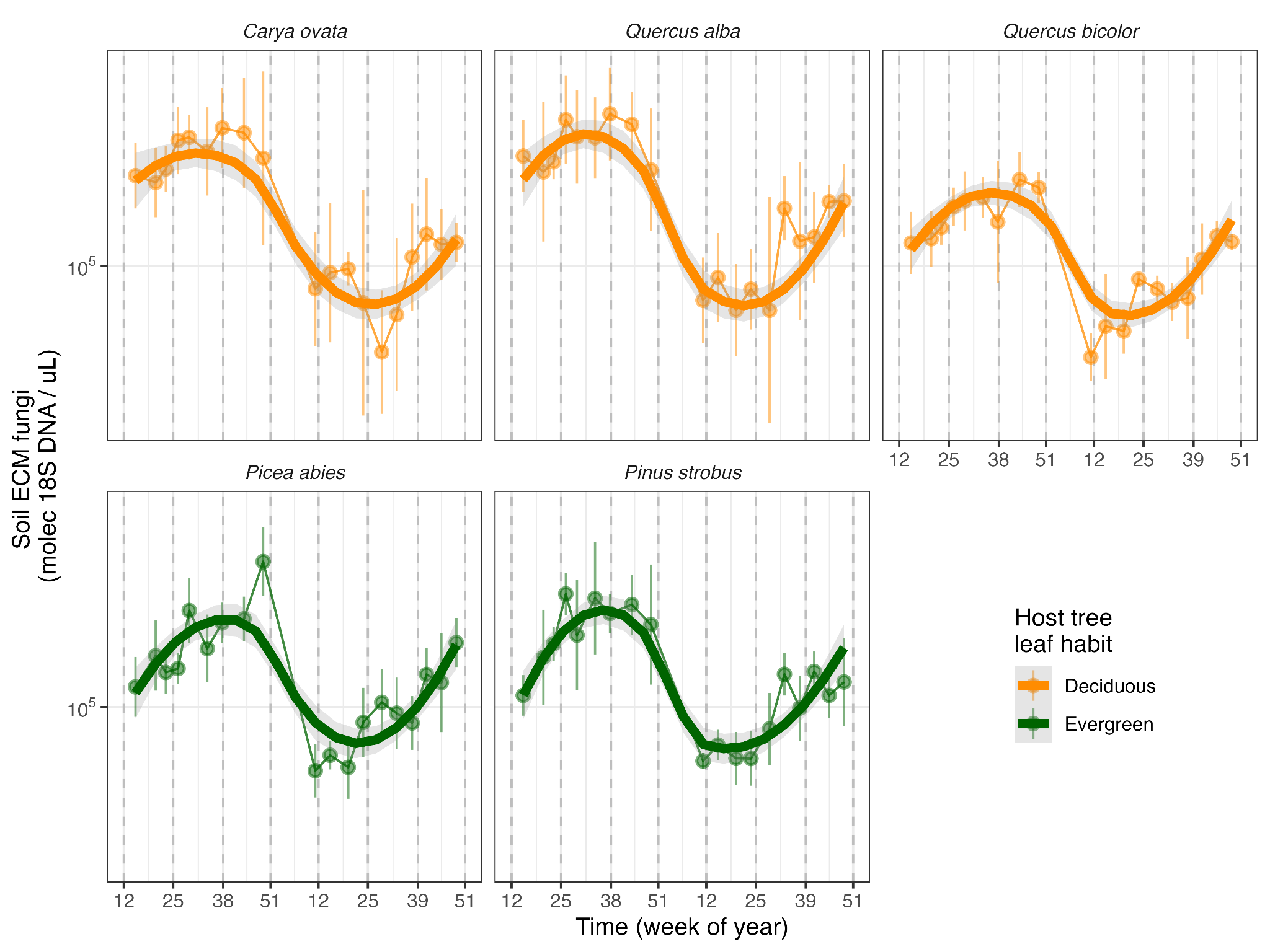

###### **Fig. S1**. Rarefied qPCR data, not normalized by year.

###### **Table S1**. Output showing generalized additive models (GAM) of quantitative soil fungal ITS2 rDNA amplification (qPCR), subset to the total proportion of sequenced DNA reads matching to ectomycorrhizal fungal lifestyles in the FungalTraits database (Põlme *et al.*, 2020), *ca.* 18%, and relativized to annual subplot maximum. Model parameter arguments include basis splines (bs) cyclical (cubic, cs) and random effect (re), and concurvity correction factor (m).

| **Component** | **Term** | **Estimate** | **Std Error** | **t-value** | **p-value** |  |
| --- | --- | --- | --- | --- | --- | --- |
| A. parametric coefficients | (Intercept) | 2,374.344 | 325.440 | 7.296 | <0.0001 | *** |
|  | **Year** | **-1.174** | **0.161** | **-7.298** | **<0.0001** | ******* |
|  | LeafHabitEvergreen | -0.003 | 0.050 | -0.053 | 0.9577 |  |
| **Component** | **Term** | **edf** | **Ref. df** | **F-value** | **p-value** |  |
| B. smooth terms | s(WEEK_unique) | 0.400 | 13.000 | 0.033 | 0.2717 |  |
|  | s(WEEK, bs = ”cs”, m = 1) | 0.000 | 5.000 | 0.000 | 0.5485 |  |
|  | **s(WEEK_unique):LeafHabitDeciduous** | **1.468** | **1.731** | **18.910** | **<0.0001** | ******* |
|  | **s(WEEK_unique):LeafHabitEvergreen** | **3.311** | **4.132** | **13.732** | **<0.0001** | ******* |
|  | ti(Temp1_7d_avg) | 1.000 | 1.000 | 0.531 | 0.4665 |  |
|  | ti(Moisture1_7d_avg) | 1.000 | 1.000 | 0.017 | 0.8960 |  |
|  | te(Temp1_7d_avg,Moisture1_7d_avg) | 1.000 | 1.000 | 2.680 | 0.1022 |  |
|  | s(SUBPLOT, bs = “re”) | 0.000 | 5.000 | 0.000 | 0.8317 |  |
| Signif. codes: 0 <= '***' < 0.001 < '**' < 0.01 < '*' < 0.05 | | | | | | |
| Adjusted R-squared: 0.220, Deviance explained 0.204 | | | | | | |
| -REML : 11.748, Scale est: 0.232, N: 562 | | | | | | |

##### Exploration type

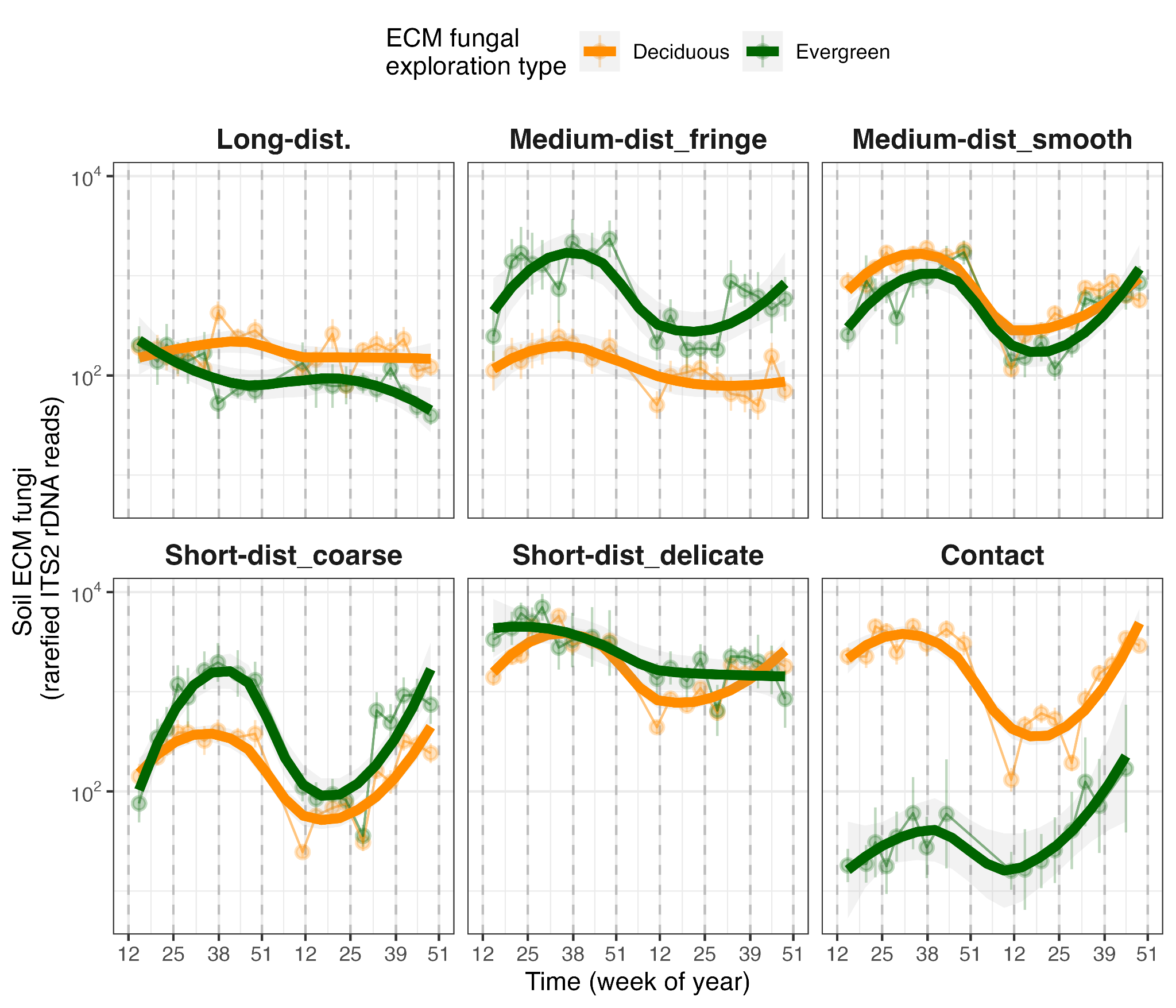

###### **Fig. S2.** Seasonality of ectomycorrhizal exploration types among five temperate monodominant forest plots, in either evergreen (dark green) or deciduous (orange) plots. Lines show smooth loess curves (*n*=6) across mean values among subplots ± 1 SE for each of 19 monthly sampling dates across the years 2023-2024 (*k*=17). Week of year number labels on x-axis indicate approximate equinox and solstice weeks.

###### **Table S2.** Output showing the generalized additive model (GAM) of ectomycorrhizal fungal exploration type abundances based on the proportion of annual maximum across subplots of rarefied eDNA abundance values, using unique week across two years and host tree species leaf habit as main predictors. Parameter arguments include basis spline (bs) as random error (“re”).

| **Component** | **Term** | **Estimate** | **Std Error** | **t-value** | **p-value** |  |
| --- | --- | --- | --- | --- | --- | --- |
| A. parametric coefficients | (Intercept) | 3,752.331 | 632.003 | 5.937 | <0.0001 | *** |
|  | **YEAR** | **-1.855** | **0.312** | **-5.940** | **<0.0001** | ******* |
|  | LeafHabitEvergreen | 0.042 | 0.041 | 1.011 | 0.3123 |  |
|  | ETMedium-dist_fringe | -0.035 | 0.059 | -0.585 | 0.5588 |  |
|  | **ETMedium-dist_smooth** | **0.455** | **0.058** | **7.885** | **<0.0001** | ******* |
|  | **ETShort-dist_coarse** | **0.150** | **0.058** | **2.576** | **0.0100** | ***** |
|  | **ETShort-dist_delicate** | **0.722** | **0.057** | **12.572** | **<0.0001** | ******* |
|  | **ETContact** | **0.552** | **0.060** | **9.225** | **<0.0001** | ******* |
| **Component** | **Term** | **edf** | **Ref. df** | **F-value** | **p-value** |  |
| B. smooth terms | **s(WEEK_unique):ETLong-dist.** | **7.073** | **8.663** | **11.312** | **<0.0001** | ******* |
|  | **s(WEEK_unique):ETMedium-dist_fringe** | **6.327** | **7.799** | **5.327** | **<0.0001** | ******* |
|  | **s(WEEK_unique):ETMedium-dist_smooth** | **1.004** | **1.008** | **14.720** | **0.0001** | ******* |
|  | **s(WEEK_unique):ETShort-dist_coarse** | **5.276** | **6.533** | **7.863** | **<0.0001** | ******* |
|  | **s(WEEK_unique):ETShort-dist_delicate** | **1.009** | **1.018** | **9.646** | **0.0019** | ****** |
|  | **s(WEEK_unique):ETContact** | **4.244** | **5.276** | **5.513** | **<0.0001** | ******* |
|  | s(SUBPLOT, bs = “re”) | 0.164 | 1.000 | 0.197 | 0.2734 |  |
|  | ti(Temp1_7d_avg) | 1.001 | 1.002 | 0.238 | 0.6260 |  |
|  | **ti(Moisture1_7d_avg)** | **7.084** | **8.257** | **2.450** | **0.0081** | ****** |
|  | te(Temp1_7d_avg,Moisture1_7d_avg) | 3.221 | 3.641 | 0.706 | 0.4116 |  |
| Adjusted R-squared: 0.170, Deviance explained 0.138 | | | | | | |
| fREML : 8698.493, Scale est: 1.391, N: 5096 | | | | | | |

##### EcM fungal genera

###### **Table S3.** Output showing the generalized additive model (GAM) of ectomycorrhizal fungal genus abundances, using week of year and host tree species leaf habit as main predictors.

| **Component** | **Term** | **Estimate** | **Std Error** | **t-value** | **p-value** |  |
| --- | --- | --- | --- | --- | --- | --- |
| A. parametric coefficients | (Intercept) | 5,691.276 | 535.789 | 10.622 | <0.0001 | *** |
|  | **Genus_*Tuber*** | **0.718** | **0.096** | **7.501** | **<0.0001** | ******* |
|  | **Genus_*Tomentella*** | **1.068** | **0.097** | **10.989** | **<0.0001** | ******* |
|  | **Genus_*Thelephora*** | **-1.266** | **0.121** | **-10.456** | **<0.0001** | ******* |
|  | **Genus_*Suillus*** | **-0.648** | **0.154** | **-4.205** | **<0.0001** | ******* |
|  | **Genus_*Sebacina*** | **0.936** | **0.116** | **8.043** | **<0.0001** | ******* |
|  | Genus_*Scleroderma* | 0.076 | 0.113 | 0.667 | 0.5046 |  |
|  | **Genus_*Russula*** | **1.845** | **0.102** | **18.089** | **<0.0001** | ******* |
|  | **Genus_*Rhizopogon*** | **-2.991** | **0.303** | **-9.863** | **<0.0001** | ******* |
|  | **Genus_*Piloderma*** | **1.004** | **0.171** | **5.870** | **<0.0001** | ******* |
|  | Genus_*Lactarius* | 0.195 | 0.142 | 1.374 | 0.1696 |  |
|  | **Genus_*Inocybe*** | **1.170** | **0.093** | **12.631** | **<0.0001** | ******* |
|  | Genus_*Hymenogaster* | 0.094 | 0.100 | 0.945 | 0.3445 |  |
|  | **Genus_*Hygrophorus*** | **1.593** | **0.127** | **12.501** | **<0.0001** | ******* |
|  | Genus_*Humaria* | 0.236 | 0.133 | 1.777 | 0.0756 | . |
|  | **Genus_*Helvella*** | **-0.519** | **0.122** | **-4.260** | **<0.0001** | ******* |
|  | **Genus_*Elaphomyces*** | **0.850** | **0.103** | **8.249** | **<0.0001** | ******* |
|  | **Genus_*Cortinarius*** | **0.413** | **0.105** | **3.936** | **0.0001** | ******* |
|  | Genus_*Clavulina* | -0.042 | 0.179 | -0.232 | 0.8167 |  |
|  | Genus_*Cenococcum* | 0.112 | 0.126 | 0.888 | 0.3746 |  |
|  | Genus_*Amanita* | 0.242 | 0.190 | 1.277 | 0.2015 |  |
|  | **LeafHabitEvergreen** | **0.512** | **0.055** | **9.353** | **<0.0001** | ******* |
|  | **YEAR** | **-2.810** | **0.265** | **-10.612** | **<0.0001** | ******* |
| **Component** | **Term** | **edf** | **Ref. df** | **F-value** | **p-value** |  |
| B. smooth terms | **s(WEEK_unique):Genus_*Wilcoxina*** | **5.130** | **5.708** | **17.629** | **<0.0001** | ******* |
|  | **s(WEEK_unique):Genus_*Tuber*** | **5.275** | **5.778** | **10.452** | **<0.0001** | ******* |
|  | **s(WEEK_unique):Genus_*Tomentella*** | **1.438** | **1.718** | **11.968** | **0.0005** | ******* |
|  | **s(WEEK_unique):Genus_*Thelephora*** | **3.564** | **4.294** | **14.213** | **<0.0001** | ******* |
|  | **s(WEEK_unique):Genus_*Suillus*** | **5.489** | **5.894** | **16.686** | **<0.0001** | ******* |
|  | **s(WEEK_unique):Genus_*Sebacina*** | **2.351** | **2.908** | **9.061** | **<0.0001** | ******* |
|  | **s(WEEK_unique):Genus_*Scleroderma*** | **5.185** | **5.739** | **25.428** | **<0.0001** | ******* |
|  | **s(WEEK_unique):Genus_*Russula*** | **2.282** | **2.803** | **6.493** | **0.0005** | ******* |
|  | **s(WEEK_unique):Genus_*Rhizopogon*** | **3.092** | **3.727** | **6.461** | **0.0001** | ******* |
|  | s(WEEK_unique):Genus_*Piloderma* | 1.000 | 1.000 | 2.087 | 0.1486 |  |
|  | **s(WEEK_unique):Genus_*Lactarius*** | **2.163** | **2.700** | **2.865** | **0.0336** | ***** |
|  | **s(WEEK_unique):Genus_*Inocybe*** | **4.874** | **5.525** | **6.422** | **<0.0001** | ******* |
|  | **s(WEEK_unique):Genus_*Hymenogaster*** | **2.384** | **2.939** | **10.068** | **<0.0001** | ******* |
|  | **s(WEEK_unique):Genus_*Hygrophorus*** | **4.336** | **5.065** | **10.762** | **<0.0001** | ******* |
|  | **s(WEEK_unique):Genus_*Humaria*** | **4.129** | **4.892** | **8.881** | **<0.0001** | ******* |
|  | **s(WEEK_unique):Genus_*Helvella*** | **4.653** | **5.326** | **9.219** | **<0.0001** | ******* |
|  | **s(WEEK_unique):Genus_*Elaphomyces*** | **4.000** | **4.756** | **9.201** | **<0.0001** | ******* |
|  | **s(WEEK_unique):Genus_*Cortinarius*** | **5.205** | **5.719** | **23.647** | **<0.0001** | ******* |
|  | **s(WEEK_unique):Genus_*Clavulina*** | **1.000** | **1.000** | **15.074** | **0.0001** | ******* |
|  | **s(WEEK_unique):Genus_*Cenococcum*** | **1.000** | **1.001** | **18.533** | **<0.0001** | ******* |
|  | **s(WEEK_unique):Genus_*Amanita*** | **2.544** | **3.015** | **6.780** | **0.0001** | ******* |
|  | ti(Temp1_7d_avg) | 1.001 | 1.001 | 0.406 | 0.5242 |  |
|  | **ti(Moisture1_7d_avg)** | **2.228** | **2.394** | **4.352** | **0.0451** | ***** |
|  | te(Temp1_7d_avg,Moisture1_7d_avg) | 7.712 | 9.254 | 1.444 | 0.1801 |  |
|  | **s(SUBPLOT)** | **4.813** | **5.000** | **18.407** | **<0.0001** | ******* |
| Adjusted R-squared: 0.145, Deviance explained 0.254 | | | | | | |
| fREML : 57985.754, Scale est: 1.000, N: 7969 | | | | | | |

##### Community composition

###### **Table S4**. Output from PERMANOVA on community tables with no term interactions.

| **Term** | **Df** | **SumOfSqs** | **R2** | **F** | **Pr(>F)** |
| --- | --- | --- | --- | --- | --- |
| **WEEK_unique** | **1** | **0.6** | **0.005** | **3.6** | **0.002***** |
| **LeafHabit** | **1** | **31.5** | **0.238** | **176.8** | **0.001***** |
| **Moisture1_7d_avg_pct** | **1** | **0.5** | **0.004** | **3.0** | **0.01**** |
| **Temp1_7d_avg** | **1** | **0.5** | **0.004** | **2.8** | **0.007***** |
| Residual | 558 | 99.5 | 0.8 |  | – |
| Total | 562 | 132.3 | 1.0 |  | – |

###### **Table S5**. Results for separate PERMDISP ordination dispersion tests testing week and leaf habit.

| **Group** | **Df** | **Sum Sq** | **Mean Sq** | **F** | **N.Perm** | **Pr(>F)** |
| --- | --- | --- | --- | --- | --- | --- |
| WEEK_unique | 18 | 0.1 | 0.0 | 0.4 | 999 | 0.99 |
| Residuals | 544 | 4.6 | 0.0 |  |  | – |

| LeafHabit | 1 | 0.0 | 0.0 | 0.0 | 999 | 0.86 |
| --- | --- | --- | --- | --- | --- | --- |
| Residuals | 561 | 4.8 | 0.0 |  |  | – |

###### **Table S6**. Output showing the RDA model of ectomycorrhizal fungal community composition, using week of year, host tree species leaf habit, soil moisture and temperature as main predictors.

| **Term** | **Df** | **Variance** | **F** | **Pr(>F)** |
| --- | --- | --- | --- | --- |
| **WEEK_unique** | **1** | **0.0** | **1.8** | **0.045*** |
| **LeafHabit** | **1** | **0.1** | **119.7** | **0.001***** |
| Moisture1_7d_avg_pct | 1 | 0.0 | 1.5 | 0.127 |
| **Temp1_7d_avg** | **1** | **0.0** | **2.4** | **0.015*** |
| WEEK_unique:LeafHabit | 1 | 0.0 | 1.3 | 0.219 |
| Moisture1_7d_avg_pct:Temp1_7d_avg | 1 | 0.0 | 0.3 | 0.999 |
| Residual | 556 | 0.5 |  | – |

###### **Fig. S3** Ordination at the species level showing intraspecific differences in RDA vector directions, and monthly sample community centroid points from evergreen (green) and deciduous (orange) plots.

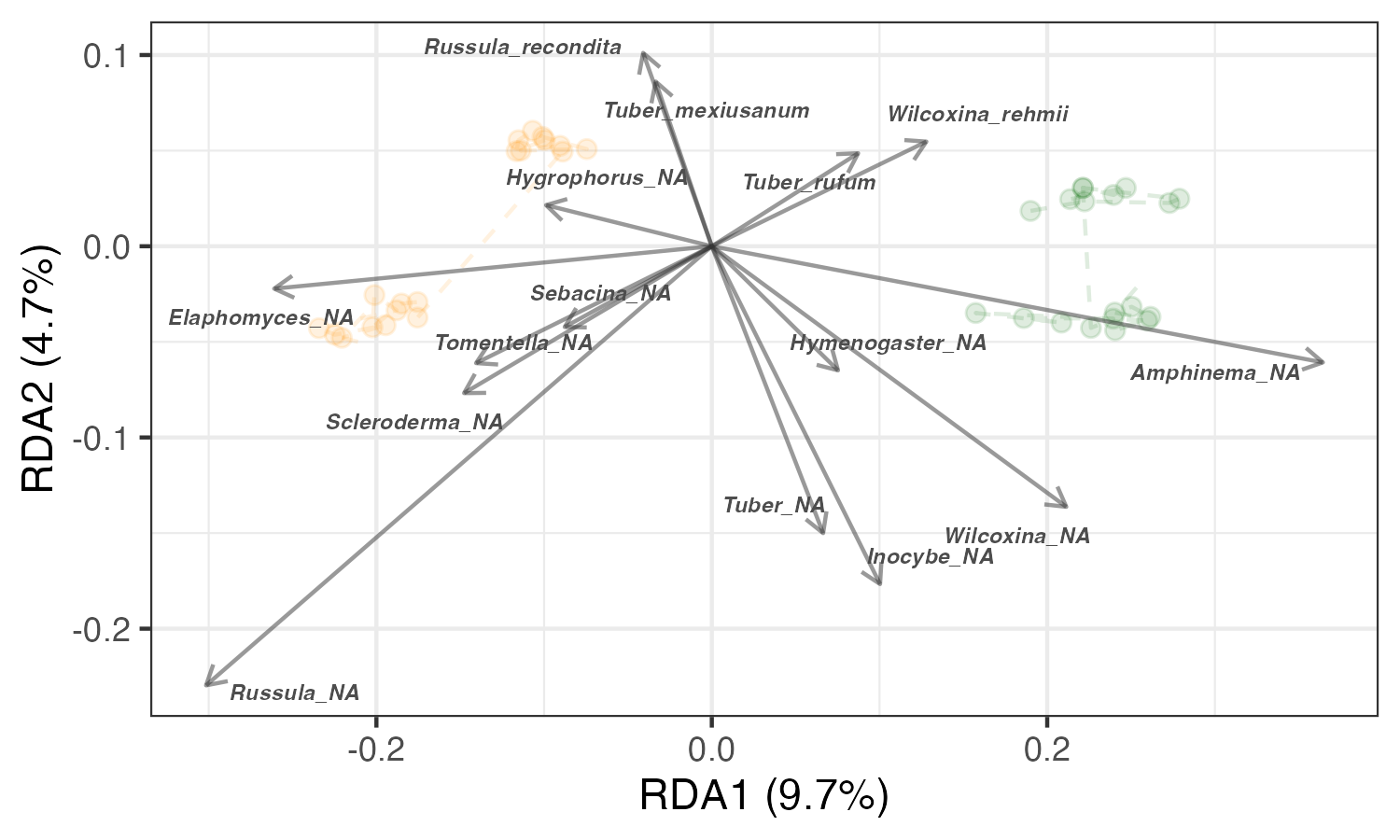

###### **Table S7** Results of PERMANOVA model on species-level community matrix.

| **Term** | **Df** | **SumOfSqs** | **R2** | **F** | **Pr(>F)** |
| --- | --- | --- | --- | --- | --- |
| **WEEK_unique** | **1** | **4.3** | **0.0** | **17.5** | **0.001** |
| **LeafHabit** | **1** | **24.2** | **0.1** | **97.3** | **0.001** |
| **Moisture1_7d_avg_pct** | **1** | **0.7** | **0.0** | **2.6** | **0.004** |
| **Temp1_7d_avg** | **1** | **0.6** | **0.0** | **2.3** | **0.006** |
| Residual | 558 | 138.9 | 0.8 |  | – |
| Total | 562 | 175.0 | 1.0 |  | – |

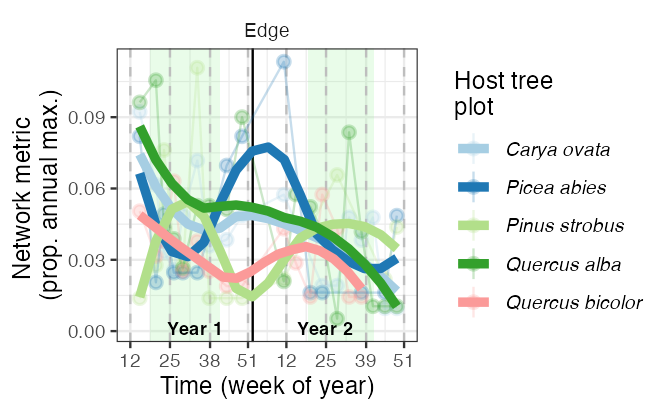

###### **Fig. S4** Dynamics of soil ectomycorrhizal fungal co-occurrence network properties, specifically number of significantly detectable edges, based on community tables at the genus level including both years of monthly samples across five monodominant temperate forest stands.
